# Reduced axonal caliber and white matter changes in a rat model of Fragile X syndrome with a deletion of a K Homology domain of *Fmr1*

**DOI:** 10.1101/864371

**Authors:** Carla E. M. Golden, Yohan Yee, Victoria X. Wang, Hala Harony-Nicolas, Patrick R. Hof, Jason P. Lerch, Joseph D. Buxbaum

**Affiliations:** Department of Psychiatry, Icahn School of Medicine at Mount Sinai, New York, NY, USA; Seaver Autism Center for Research and Treatment, Icahn School of Medicine at Mount Sinai, New York, NY, USA; Department of Medical Biophysics, University of Toronto, Toronto, Ontario, Canada; Mouse Imaging Centre, The Hospital for Sick Children, Toronto, Ontario, Canada; BioMedical Engineering and Imaging Institute, Icahn School of Medicine at Mount Sinai, New York, NY, USA; Nash Family Department of Neuroscience, Icahn School of Medicine at Mount Sinai, New York, NY, USA; Friedman Brain Institute, Icahn School of Medicine at Mount Sinai, New York, NY, USA; Mindich Child Health and Development Institute, Icahn School of Medicine at Mount Sinai, New York, NY, USA; Wellcome Centre for Integrative Neuroimaging, University of Oxford, Oxford, UK; Department of Genetics and Genomic Sciences, Icahn School of Medicine at Mount Sinai, New York, NY, USA

## Abstract

Fragile X syndrome (FXS) is a neurodevelopmental disorder that is caused by mutations in the *FMR1* gene that are known to cause neuroanatomical alterations. The morphological underpinnings of these alterations have not been elucidated. Furthermore, while alterations have been identified in both male and female individuals, neuroanatomy in female rodent models has not been assessed. We identified structural differences in regions that are also altered in FXS in male and female rat models, including the splenium of the corpus callosum. Interestingly, different sets of regions were disrupted in male and female rat models and, remarkably, male rats had higher brain-wide diffusion than female rats overall. We found reduced axonal caliber in the splenium, offering a mechanism for its structural changes. Our results provide insight into which brain regions are vulnerable to a loss of *Fmr1* expression and suggest a potential mechanism for how its loss causes white matter dysfunction in FXS.

## Introduction

Fragile X syndrome (FXS) is a monogenic disorder caused by mutations in the *FMR1* gene, which encodes fragile X mental retardation protein (FMRP). It is the leading monogenic cause of autism spectrum disorder (ASD), the most frequent known form of inherited intellectual disability (ID), is often comorbid with attention-deficit/hyperactivity disorder (ADHD), and can cause sensory hyperarousal [1, 2]. With the use of magnetic resonance imaging (MRI), alterations in brain structure have been identified in both gray and white matter regions in individuals with FXS and associated with aberrant cognitive phenotypes. However, the integrity of brain regions and the white matter that connects them has not been directly assessed in a rat model of FXS. The regions that should be given priority in studies of FXS rat models have therefore not been elucidated. Furthermore, the mechanism by which a lack of FMRP causes structural alterations is poorly understood and can only be studied in rodent models of FXS.

In humans with FXS, impairments that change across development have been identified in frontal and striatal regions and the white matter that connects them [3]. It is hypothesized that these alterations underlie the reported deficits in attention in FXS. However, other regions and white matter tracts that are also implicated in attention show anatomical alterations in FXS, such as the thalamus, internal capsule, and the splenium of the corpus callosum [4]. Structural deficits in white matter have also been identified in *Fmr1* knockout (KO) mice and resemble those seen in individuals with FXS [5–7]. Importantly, this includes regions known to be recruited during visuospatial attention, including the superior colliculus, and the tracts that connect them, including the splenium of the corpus callosum and the white matter of the medial prefrontal cortex. There are also deficits in the degree to which networks containing spatially disparate regions fire together while the individual is at rest, called functional connectivity, in individuals with FXS and *Fmr1* KO mice [8, 9]. One example of this is hypoconnectivity in the somatosensory network of *Fmr1* KO mice, again suggesting an anatomical link between decreased Fmrp expression and deficits in sensory perception [9]. We predict that similar regions will be impaired in a rat model of FXS.

FMRP could play a role in structural integrity through its repression of protein translation of mRNA targets [10]. In FXS, without FMRP, the protein expression of these mRNA targets is elevated because their translation is no longer repressed. Many of the mRNA targets of Fmrp encode key presynaptic and postsynaptic proteins that affect synapse development, including cytoskeleton scaffolding and remodeling, as well as transcripts involved in the development of myelin [1]. Dysregulation of their expression in the absence of FMRP is thought to contribute to the heightened levels of long, immature dendritic spines and increased spine density that is commonly seen in the pyramidal neurons of individuals with FXS and animal models [1]. Therefore, these alterations in neurons and oligodendrocytes could influence the brain’s functional anatomy, however, a mechanism by which this might occur has not yet been elucidated. Thus, it is pertinent to perform an unbiased study of structural integrity in rats that screens across regions.

We recently identified a deficit in sustained attention in both sexes of a rat model of FXS that has a deletion of an mRNA-binding domain, K-homology 1 (KH1), the *Fmr1-^Δ^exon 8* rat [11]. Instead of Fmrp, the *Fmr1-^Δ^exon 8* rat expresses Fmrp-^Δ^KH1 at significantly low levels. Auditory deficits have also been modeled in this rat model with impaired cortical representation of speech sounds [12]. We chose to assess neuroanatomical integrity in this *Fmr1-^Δ^exon 8* rat model of FXS. We used structural MRI to measure regional volumes and tissue densities, diffusion tensor imaging (DTI) to assess white matter integrity, and electron microscopy to uncover deficits in ultrastructure that could reflect functional perturbations. Both male and female *Fmr1-^Δ^exon 8* rats were included in these studies to test for an effect of sex and for a dose-dependent effect of Fmrp-^Δ^KH1 in a rat that is heterozygous in expression for *Fmr1-^Δ^exon 8*, a phenomenon only possible in females.

## Materials and Methods

### Experimental Design

The objective of this controlled laboratory experiment was to use an unbiased approach to identify brain regions that are affected by a loss of *Fmr1* expression and then determine the mechanism that could underlie this change. We hypothesized that brain regions that are affected in individuals with FXS would also be altered in the *Fmr1-^Δ^exon 8* rat model of FXS. Once we found that diffusion in the splenium of the corpus callosum was altered, we hypothesized that this could be due to alterations in axon integrity. Sample sizes used in the MRI experiments were determined based on a previous study of a rodent model of autism [13] and sample sizes for electron microscopy mirrored similar studies. Sex was considered a variable and its effect was reported where meaningful. Analysis of the MRI data was replicated with two different analytical pipelines and electron microscopy was replicated in two separate cohorts, each constituting a sample size of three. The degree to which the MRI results were validated in the second analysis are detailed in the Results section and the findings from the electron microscopy experiments were substantiated in the second cohort. During image acquisition, manual quality control of the MRI segmentations, and tracing of the electron microscopy images, the experimenters were blind to genotype. Furthermore, each manually performed operation was validated across two experimenters to limit subjectivity. Each experimental animal was scanned in the MRI the same day as a same-sex control animal. Images with obvious artifacts or masks that did not align to the image were excluded before the statistical analyses commenced. Individual data points that were outside 1.5 times the interquartile range were omitted from the analysis. No outliers were included in the results reported.

### Generation of the *Fmr1-^Δ^exon 8* rat model

The *Fmr1-^Δ^exon 8* rat model was generated using zinc finger nucleases (ZFNs) in the outbred Sprague-Dawley background. The design and cloning of the ZFN, as well as the embryonic microinjection and screening for positive founder rats were performed by Horizon Labs (Boyertown, PA USA) as previously described [14]. The best performing ZFN pair targeting the CATGAACAGTTTATCgtacgaGAAGATCTGATGGGT sequence, located between 18686bp–18721bp in the *Fmr1* gene (NCBI reference sequence NC_005120.4), was used for embryo microinjection. Positive Sprague-Dawley founder animals with a deletion in the *Fmr1* gene were mated to produce F1 breeding pairs. Polymerase chain reaction amplification at the target sites followed by sequencing analysis revealed the exact deletion of 122 bp at the junction of intron 7 and exon 8 (between 18588bp-18709bp).

### Animal breeding, care, and husbandry

This study used age-matched male and female littermate rats. To produce both male genotypes (*Fmr1-^Δ^exon 8 ^+/y^ (WT) and Fmr1-^Δ^exon 8^-/y^*) and all three female genotypes (*Fmr1-^Δ^exon 8^+/+^ (WT), Fmr1-^Δ^exon 8^+/-^, and Fmr1-^Δ^exon 8^-/-^*), we set up two pairs of breeders: *WT* male x *Fmr1-^Δ^exon 8^+/-^ and Fmr1-^Δ^exon 8^-/y^* x *Fmr1-^Δ^exon 8^+/-^*. All five groups were used for the MRI experiments and WT and *Fmr1-^Δ^exon 8^-/y^* male rats were used for the electron microscopy experiments. All rats were kept under veterinary supervision in a 12 h reverse light/dark cycle at 22±2°C. Animals were pair-caged with food and water available *ad libitum*. All animal procedures were approved by the Institutional Animal Care and Use Committee at the Icahn School of Medicine at Mount Sinai.

### MRI

All imaging was performed by the BioMedical Molecular Imaging Institute using a Bruker Biospec 70/30 7 T scanner with a B-GA12S gradient insert (gradient strength 440 mT/m and slew rate 3444 T/m/s). A Bruker 4 Channel rat brain phased array was used for all data acquisition in conjunction with a Bruker volume transmit 86-cm coil. All rats (N = 15/group) were imaged on a heated bed and respiration was monitored continuously until the end of the scan. The animal was anesthetized using isoflurane anesthesia (3% induction and 1.5% maintenance). After a three-plane localizer, a field map was acquired and the rat brain was shimmed using Mapshim software. A DTI protocol was acquired with a Pulsed Gradient Spin Echo – Echo-planar imaging (EPI) sequence with the following parameters: repetition time (TR) = 5000ms, echo time (TE) = 22.6 ms, 4 segments, 30 gradient directions with b-value = 1000 s/mm^2^ and 5 B0’s, field of view (FOV) = 25 mm, Matrix = 128×128, slice thickness = 1 mm, skip = 0, 6 averages, total acquisition time = 1 hr. The voxel size was 0.195 x 0.195 x 1 mm^3^ (1000 µm-thick). A high resolution T2 anatomical scan was obtained with a 3D Rapid Acquisition with Relaxation Enhancement (RARE) sequence with a RARE factor of 8, TR = 777 ms, effective TE = 52 ms, FOV = 30 mm x 27.25 mm x 30 mm, matrix size 256 x 256 x 128. The voxel size was 0.117 x 0.117 x 0.234 mm^3^ (234 µm-thick).

### MRI region-based analytical pipeline with manual editing

A magnetic resonance imaging processing pipeline was used to perform semi-automated nonbiased brain segmentation, while blinded to genotype (N = 15/group) [15]. The pipeline is composed of six major steps: rigid registration of images to each other, generation of a whole-brain mask for each image, averaging of all images, creation of a whole-brain mask for this averaged image, segmentation of the average mask by regions of interest (ROIs), parcellation propagation of the segmented mask to individual subjects, and ROI-based statistics for the individual images. The deformation necessary to warp each subject’s image to the average was used to calculate the volume of the ROIs. After each mask was generated, it was improved manually in ITK-SNAP (www.itksnap.org). The segmentation into ROIs was determined by a template that was previously hand-segmented into 32 brain regions, listed in **Supplementary Table 1**.

Segmented masks for the individual images that did not closely match the segmented average mask (one *Fmr1-^Δ^exon 8 ^-/-^* T2 mask, two female WT T2 masks, and one male WT DTI mask) and individual data point outliers, defined as being outside 1.5 times the interquartile range, were excluded from the analysis. Males and females were analyzed in two separate pipelines because the difference in their brain size would skew the averaging step. The whole-brain masks were used to determine whole-brain measures. Mean voxel intensity for each ROI and across the whole-brain was measured in both the T2 and DTI images and the volume of each ROI and whole-brain was calculated from the T2 images. The olfactory bulb, cerebellum, and brainstem were not included in the DTI analysis because these images did not capture the entirety of these regions. Furthermore, regions without any known white matter were excluded from the DTI analysis, including the aqueduct, periaqueductal gray (PAG), and third, fourth, and lateral ventricles. In the analysis of the male data, for each independent variable, if the distribution was nonparametric according to the Shapiro-Wilk’s test, a Mann-Whitney U test was administered and if the distribution was normally distributed, a two-way ANOVA was applied. In the analysis of the female data, if the distribution was nonparametric, a Kruskal-Wallis test was administered, which was followed by a Dunn test to compare the individual means and an adjustment of the *p*-values with the False Discovery Rate method, and if the data was parametric, an ANOVA was applied and followed by a post-hoc Tukey HSD test that compared the pairs of means and adjusted the *p*-values to account for the additional comparisons. Due to the fact that many comparisons were made across ROIs, the output was then assessed for the ability to survive a correction for multiple comparisons with a Bonferroni Correction, the most conservative method of this nature. Genotype was the only between groups factor. Custom scripts written in the R statistical programming environment were used for the statistical analysis (R Development Core Team, 2006).

### Automated MRI region-based and voxel-based analytical pipeline

T2 images were aligned together in an iterative registration procedure using the PydPiper framework that included a series of linear and nonlinear alignment steps, as previously described [16]. Briefly, images were first linearly aligned (6 degrees of freedom: rotations and translations) to a model template, so that all images were roughly in the same space and roughly in the same orientation. These images were then linearly aligned (12 degrees of freedom: rotations, translations, scaling and shearing) to each other in a pairwise manner; each image was resampled with its average transformation to the other images. The linearly transformed resampled images were then averaged to create a linear (12-parameter) template. After creation of a linear template, images were then nonlinearly aligned with Advanced Normalization Tools (ANTs) [17] (www.picsl.upenn.edu/software/ants) to this template and averaged to create a new updated template. Nonlinear alignment and averaging were repeated for a total of three iterations. The output of this pipeline included a final study-specific nonlinear average template, transformations mapping each image to the template, and voxel-wise Jacobian determinants corresponding to each transformation that represent the extent to which each image locally deformed to match the template. To identify voxels that were altered in volume, a *p*-value significance was set to an false discovery rate (FDR)-corrected *p*-value < 0.1. We controlled for the FDR instead of applying the Bonferroni method in the analysis of the voxel-wise data because the dataset was much larger. The Bonferroni method could therefore yield false-negatives.

DTI metrics were extracted for each subject from the raw imaging data with DTIFit (Functional Magnetic Resonance Imaging of the Brain Software Library). The raw T2 (no diffusion weighting, “S0”) images were registered together using the same pipeline described above. This DTI S0 template was further aligned to the T2 template using ANTs, and concatenated transformations mapping each raw image to the T2 template were computed. The remaining images (fractional anisotropy (FA), mean diffusion (MD), axial diffusion (AD), radial diffusion (RD)) were resampled to the T2 template with their respective concatenated transforms to allow both voxel-wise and ROI analyses.

We used DTI to evaluate white matter specifically, as it is affected in individuals with FXS, with AD, RD, MD, and total FA as measures of possible changes. FA is enhanced when parallel diffusivity is facilitated and/or perpendicular diffusivity is restricted, RD is increased following myelin damage, and AD is reduced after axonal damage [18]. Furthermore, MD may reflect pathology if it is increased in white matter.

The template that was used for the T2 semi-automated analysis was aligned to the T2* template using ANTs in order to segment it into ROIs (see **Supplementary Table 1**) and resampled to these DTI templates. For T2, volumes for each subject were computed by summing the voxel volumes under each ROI, weighted by the Jacobian determinants. This was done using the RMINC software library in R (https://github.com/Mouse-Imaging-Centre/RMINC). Mean volume for each ROI and each DTI metric were computed in T2* space using RMINC. Whole brain values for the DTI metrics were calculated by summing the average metrics per ROI, weighted by volume of each ROI. The regional volume of the PAG was computed using the Waxholm Space Atlas (NITRC), a better estimate of this region, once volumetric changes in voxels within it were identified.

For this analysis, *Fmr1-^Δ^exon 8^-/-^* and *Fmr1-^Δ^exon 8^-/y^* rats were compared to their WT male and female littermates (N = 15/group) so that the effect of sex could be examined. Data points that were outside 1.5 times the interquartile range were determined to be outliers and removed from the analysis. Statistical analysis began with the assessment of normality. Non-parametric data were evaluated with a linear model (LM) and parametric data was assessed with a two-way ANOVA, both with genotype and sex as factors. A Tukey HSD post hoc test was administered if there was a main effect of genotype or sex to correct for post hoc multiple comparisons. The output was corrected for multiple comparisons with a Bonferroni Correction, just as in the semi-automated ROI-based analysis.

### Electron Microscopy

Preparation for electron microscopy was performed by the Icahn School of Medicine’s Microscopy CoRE using protocols optimized to study the ultrastructure of nervous tissue. Male rats (WT: N = 7, *Fmr1-^Δ^exon 8^-/y^*: N = 6) were anesthetized and perfused using a peristaltic pump at a flow rate of 35 ml/min with 1% paraformaldehyde/phosphate buffered saline (PBS), pH 7.2, and immediately followed with 2% paraformaldehyde and 2% glutaraldehyde/PBS, pH 7.2, at the same flow rate for an additional 10-12 min. The brain was removed and placed in immersion fixation (same as above) to be postfixed for a minimum of one week at 4 °C. Fixed brains were sectioned using a Leica VT1000S vibratome (Leica Biosystems, Buffalo Grove, IL) and coronal slices (325 µm) containing the splenium of the corpus callosum were removed and embedded in EPON resin (Electron microscopy Sciences (EMS), Hatfield, PA). Briefly, sections were rinsed in 0.1 M sodium cacodylate buffer (EMS), fixed with 1% osmium tetroxide followed with 2% uranyl acetate, dehydrated through ascending ethanol series (beginning with 25% up to 100%) and infiltrated with propylene oxide (EMS) and then EPON resin (EMS). Sections were transferred to beem capsules and heat-polymerized at 60 °C for 72 h. Semithin sections (0.5 and 1 µm) were obtained using a Leica UC7 ultramicrotome, counterstained with 1% Toluidine Blue, coverslipped and viewed under a light microscope to identify and secure the region of interest. Ultrathin sections (80 nm) were collected on copper 300 mesh grids (EMS) using a Coat-Quick adhesive pen (EMS), and serial sections were collected on carbon-coated slot grids (EMS). Sections were counter-stained with 1% uranyl acetate followed with lead citrate.

Sections were then imaged on an Hitachi 7000 electron microscope (Hitachi High-Technologies, Tokyo, Japan) using an advantage CCD camera (Advanced Microscopy Techniques, Danvers, MA). Ten areas from within the region of the splenium that contained cross sections of axons were chosen randomly to be imaged per section at 15,000 x magnification. Images were adjusted for brightness and contrast using Adobe Photoshop CS4 11.0.1. Using ImageJ (www.imagej.nih.gov), 200 myelinated and 200 unmyelinated axons were traced from each sample. The caliber of the myelinated and unmyelinated axons was equal to the area within the trace of the cell walls. The thickness of the myelin was calculated by subtracting the traces of the myelinated axons from traces that included the exterior of the myelin (excluding perfusion artifacts). The number of myelinated axons per image was determined by counting by hand. The g-ratio was calculated as the caliber of each myelinated axon divided by its caliber plus the thickness of its myelin. The interaction between genotype and caliber was assessed. The amount of the total area in the image that was covered by the myelinated axons was calculated per image by multiplying the number of myelinated axons by the average caliber of the myelinated axons in the image and then comparing the result as a percentage of the total area of the image (58,253,186 nm^2^). Two cohorts of littermates were assessed where the first cohort had three WT and three *Fmr1-^Δ^exon 8^-/y^* rats and the second cohort had four WT and three *Fmr1-^Δ^exon 8^-/y^* rats. A Kolmogorov–Smirnov test was used to test for the distance between the empirical distribution functions of axon caliber. Data points that were outside 1.5 times the interquartile range for each rat were removed and a linear mixed model where the image variable was nested in the rat ID which was nested in the cohort number in order to determine the effect of genotype was computed using custom scripts written in the R statistical programming environment (R Development Core Team, 2006).

## Results

### Brain volumes are altered in *Fmr1-^Δ^exon 8* rats

Structural T2-weighted MRI was used to assess both volume and white versus gray matter density in male and female *Fmr1-^Δ^exon 8* rats and their WT littermates. The brains were divided into ROIs to determine whether these measures in the whole brain or in 32 ROIs (see **Supplementary Table 1**) were affected by genotype. In the region-based analysis, ROIs were first determined based on a template (see **Methods**) in an averaged image that was calculated from all subjects. This method of segmentation was then applied to each individual image. Because male rodents have larger brains than female rodents [19] due to increased body size, we decided to analyze the data two ways: one with the males and females separated and one with them combined. We will first describe the results of the analysis where they were split by sex.

There were no significant differences in whole-brain volume between the *Fmr1-^Δ^exon 8^-/y^* rats and their sex-matched WT littermates (**Supplementary Figure 1A;** ANOVA, *p* = 0.72). Additionally, there were no differences in mean tissue density across the whole brain and in individual ROIs, suggesting an equal distribution of gray and white matter in *Fmr1-^Δ^exon 8^-/y^* rats and their male WT littermates (**Supplementary Figure 1A;** ANOVA, *p* = 0.63). Furthermore, the volume across most ROIs did not differ by genotype and no regions differed by genotype when we normalized the volume of each ROI to the volume of the whole brain. The only metric that was altered was the absolute volume of the superior colliculus, which had a significant nominal *p*-value that did not survive a Bonferroni correction for multiple comparisons, but was increased in the *Fmr1-^Δ^exon 8^-/y^* rats (**Supplementary Figure 1B;** ANOVA, *p* = 0.011). Pertinent to this study, the superior colliculus is implicated in visuospatial attention across species [20], most specifically in spatial orienting.

Similar to the analysis of the male rats, there was no significant effect of genotype on whole-brain volume or density in the analysis of the female WT, *Fmr1-^Δ^exon 8^-/-^*, and *Fmr1-^Δ^exon 8^-/+^* littermates (**Figure 1A;** ANOVA, *p* = 0.62 and *p* = 0.55, respectively). Furthermore, none of the nominal *p*-values in any of the T2 analyses survived a Bonferroni correction for multiple comparisons. However, some of the nominal *p*-values showed that there were ROIs that differed in absolute volume. There was a significant effect of genotype on the volume of the genu of the corpus callosum (Kruskal-Wallis, *p* = 0.047) and the olfactory bulb (Kruskal-Wallis, *p* = 0.01) (**Figure 1A and B)**. The genu of the corpus callosum and the olfactory bulb were significantly larger in the *Fmr1-^Δ^exon 8^-/+^* rats compared to the female WT rats (**Figure 1B;** Dunn’s test, *p adj.* = 0.02 and *p adj.*= 0.023, respectively). Next, we examined the relative volume of each ROI by calculating its percentage of each rat’s total brain volume. We found that the genu was again increased in *Fmr1-^Δ^exon 8^-/+^* rats compared to female WT littermates (**Figure 1A and C;** ANOVA followed by Tukey HSD, *p adj.* = 0.03). Lastly, the hypothalamus was reduced in relative volume in the *Fmr1-^Δ^exon 8^-/-^* compared to WT female littermates (**Figure 1A and C;** Kruskal-Wallis followed by Dunn’s test, *p adj.* = 0.02).

**Figure 1.**
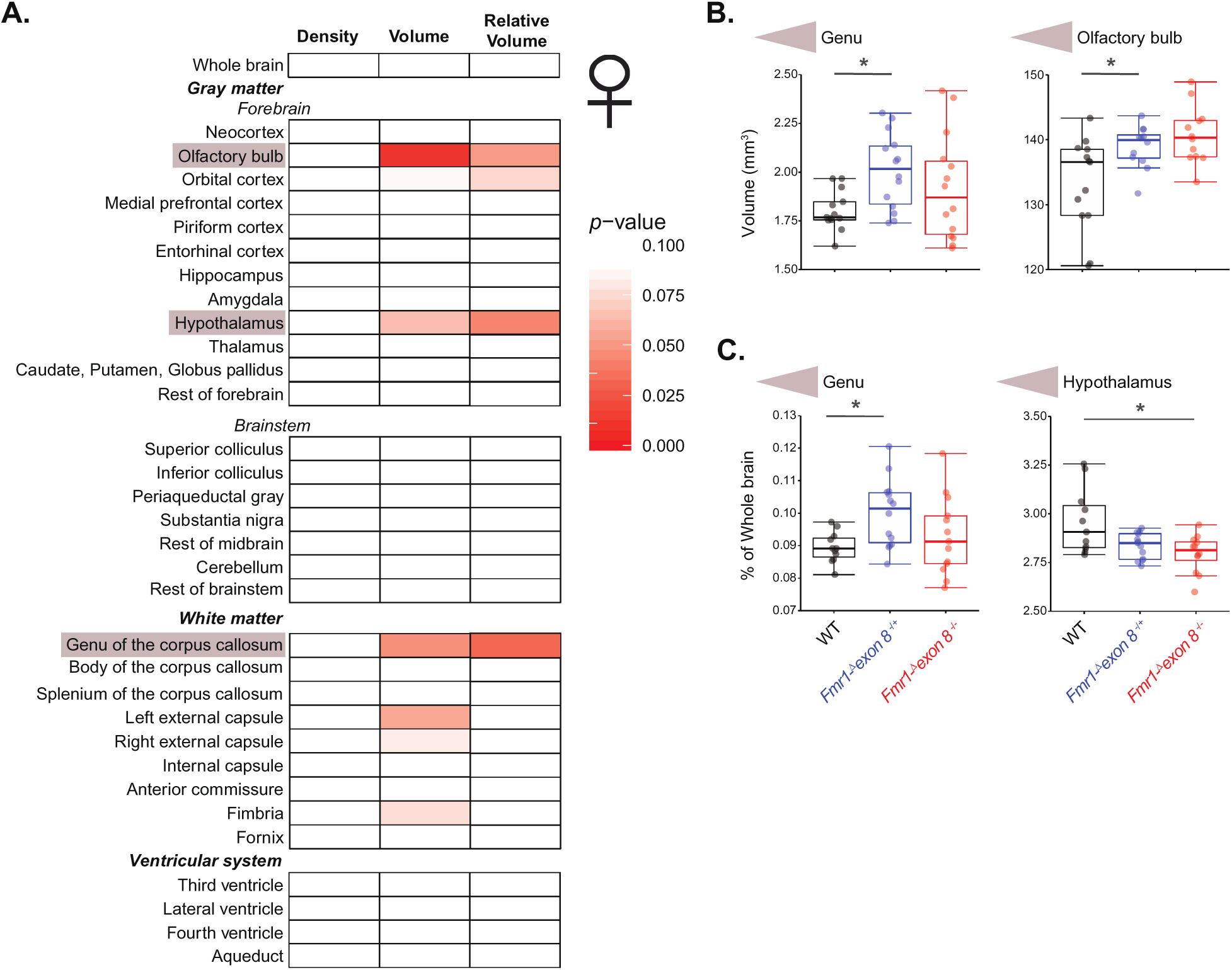
Tissue density and absolute and relative volume of ROIs in *Fmr1-^Δ^exon 8^-/-^* rats, *Fmr1-^Δ^exon 8^-/+^* rats, and female WT littermates. **(A)** Heatmap of *p*-values from ANOVA and Kruskal-Wallis tests for an effect of genotype on tissue density and absolute and relative volume in T2 images where an increase in red denotes increased significance. **(B)** Boxplots of the regions that significantly changed in absolute volume, the genu and olfactory bulb. **(C)** Boxplots of the regions that significantly changed in relative volume compared to the whole brain, the genu and hypothalamus. Significance bars represent pair-wise comparisons from either a Tukey HSD or Dunn’s test, (WT: N = 13; *Fmr1-^Δ^exon 8^-/+^*: N = 15; *Fmr1-^Δ^exon 8^-/-^*: N = 14), \**p* < 0.05.

### Volumetric changes are validated when males and females are combined

We next validated our findings with a second, fully automated pipeline that used the same method of segmentation (see **Supplementary Table 1** for ROIs) and combined males and females. All brains were first registered together and then ROIs were determined for each subject individually, thus allowing us to directly compare males and females. *Fmr1-^Δ^exon 8^-/+^* rats were excluded so that there was an equal number of genotypes represented per sex (WT and *Fmr1-^Δ^exon 8* rats). This analysis confirmed our hypothesis that male brains would be larger than female brains (**Supplementary Figure 2A and B**; LM followed by Tukey HSD, *p adj.* < 1.0 x 10^-20^) with a nominal *p*-value that survives a Bonferroni correction and showed that most brain regions increase in volume. Furthermore, it validated our finding that the absolute volume of the superior colliculus was increased in *Fmr1-^Δ^exon 8* rats, this time in both male and female *Fmr1-^Δ^exon 8* rats and with a *p*-value that survives a Bonferroni correction (**Figure 2A and B**; LM followed by Tukey HSD, *p adj.* = 4.67 x 10^-5^). It also identified an increase in the relative volume of the superior colliculus in male and female *Fmr1-^Δ^exon 8* rats, which also survived a Bonferroni correction (**Figure 2A and B**; LM followed by Tukey HSD, *p adj.* = 1.59 x 10^-10^). There was a significant effect of sex on absolute and relative volume of the superior colliculus (**Figure 2A and B**; LM followed by Tukey HSD, *p* = 4.4 x 10^-13^ and *p* = 5.08 x 10^-8^, respectively). Additionally, the relative volume of the lateral ventricle increased and the relative volume of the fourth ventricle decreased in *Fmr1-^Δ^exon 8* rats (**Figure 2A and B**; two-way ANOVA followed by Tukey HSD, *p adj.* = 2.01 x 10^-7^ and *p adj.* = 2.11 x 10^-4^, respectively). The superior colliculus and fourth ventricle were larger in the male rats (**Supplementary Figure 2A**).

**Figure 2.**
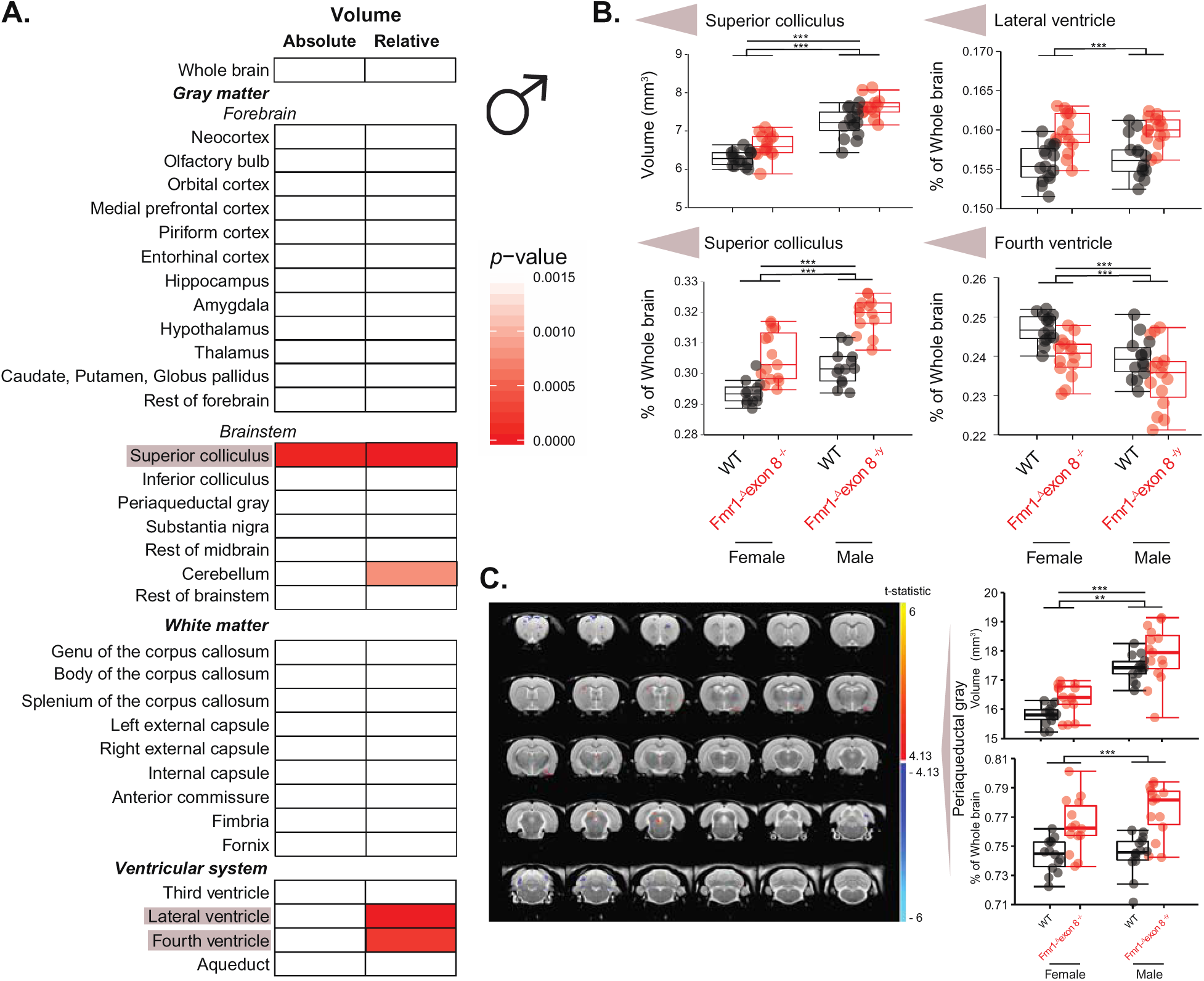
Absolute and relative brain region volumes of WT and *Fmr1-^Δ^exon 8* littermates. **(A)** Heatmap of *p*-values from ANOVA and LM tests for an effect of genotype on absolute and relative volume in T2 images where an increase in red denotes increased Bonferroni-corrected significance. **(B)** Boxplots of the regions that significantly changed in absolute and relative volumes, the superior colliculus and lateral and fourth ventricles. **(C)** Map of the t-statistic across the brain showing the voxels that are increased in volume in the *Fmr1-^Δ^exon* rats compared to WT rats. Boxplots of mean **(E)** absolute and **(E)** relative volume of a region that contained voxels that were significantly altered, the PAG, in *Fmr1-^Δ^exon* rats. Significance bars represent pair-wise comparisons from a Tukey HSD test, (N = 15/genotype), \*\*\**p* < 0.001, \*\**p* < 0.01.

### Voxel-based analysis identifies further volumetric changes

Because this pipeline transforms each image to a study-based average, we were additionally able to test for local effects that would not otherwise be captured at the structural level with a voxel-based approach. The PAG contained many voxels that were affected by genotype (**Figure 2C**; *p*FDR < 0.1). After applying region segmentation using the Waxholm Space Atlas, we found that there was a main effect of genotype that survived a Bonferroni correction (**Figure 2C**; two-way ANOVA followed by Tukey HSD, *p adj.* = 2 x 10^-7^) on relative volume of the PAG where the *Fmr1-^Δ^exon 8* rats had an increased volume. There was no effect of sex (**Figure 2C**; *p* = 0.267). The absolute volume of the PAG was also significantly increased in *Fmr1-^Δ^exon 8* compared to WT rats and in male rats compared to female rats (**Figure 2C;** LM followed by Tukey HSD, *p adj.* = 0.002 and *p adj.* = 7.46 x 10^-13^, respectively); however, the effect of genotype did not survive a Bonferroni correction and the effect of sex was likely due to the overall increase in brain size in males.

### Diffusion is altered in *Fmr1-^Δ^exon 8* rats

We used DTI to evaluate white matter specifically, as it is affected in individuals with FXS, with axial, radial, and mean diffusion (AD, RD, and MD, respectively) and total FA as measures of possible changes. FA is enhanced when parallel diffusivity is facilitated and/or perpendicular diffusivity is restricted, RD is increased following myelin damage, and AD is reduced after axonal damage [18]. Furthermore, MD may reflect pathology if it is increased in white matter.

Similar to the T2 region-based analysis, we first analyzed the male and female rats separately and assessed both the whole-brain and the 32 brain ROIs (see **Supplementary Table 1**). When we compared *Fmr1-^Δ^exon 8^-/y^* rats to their male WT littermates, we found that the nominal *p*-values for the DTI metrics of multiple ROIs were significant, though only a few survived a Bonferroni correction. Remarkably, RD was increased across the whole brain (**Figure 3A and B**; ANOVA, *p* = 0.043). This could be due to RD being increased in many forebrain regions, including the largest region of the brain in our analysis, the neocortex (**Figure 3A and C**; ANOVA, *p* = 0.015). RD also increased in the piriform cortex, hypothalamus, and thalamus (**Figure 3A and C**; ANOVA, *p* = 0.0009, *p* = 0.0017, and *p* = 0.0047, respectively), as well as the caudate, putamen, and globus pallidus (**Figure 3A and C**; ANOVA, *p* = 0.032), which are known to be implicated in impulsivity [21] and were grouped together as one region in our analysis. The nominal *p*-values associated with the increase in RD in the piriform cortex and hypothalamus survived a Bonferroni correction. Complementary to this increase in RD, there were also increases in MD in the forebrain in the piriform cortex, hypothalamus, and thalamus (**Figure 3A and C**; ANOVA, *p* = 0.00012, *p* = 0.0056, and *p* = 0.02, respectively). The nominal *p*-value associated with the increase in MD in the piriform cortex survived a Bonferroni correction. Interestingly, AD, though often inversely proportional to RD and MD, also increased in many forebrain regions. AD was higher in the neocortex, piriform cortex, hypothalamus, and thalamus (**Figure 3A and C**; ANOVA, *p* = 0.044, *p* = 0.0067, *p* = 0.039, and *p* = 0.047, respectively). While there were no alterations in these indices in the brainstem regions, there were changes in white matter pathways, as we predicted. All three diffusion indices, AD, MD, and RD, increased in the splenium of the corpus callosum (**Figure 3A and C**; ANOVA, *p* = 0.0052, *p* = 0.003, and *p* = 0.0035). The nominal *p*-values for the increase in MD and RD in the splenium trended towards significance after being corrected for multiple comparisons. MD and RD were enhanced in the internal capsule (**Figure 3A and C**; ANOVA, *p* = 0.019 and *p* = 0.0065) and AD and MD were greater in the fimbria (**Figure 3A and C**; ANOVA, *p* = 0.029 and *p* = 0.028). This whole-brain increase in RD and these additional deficits to white matter regions suggest a potential brain-wide deficit in axonal projections.

**Figure 3.**
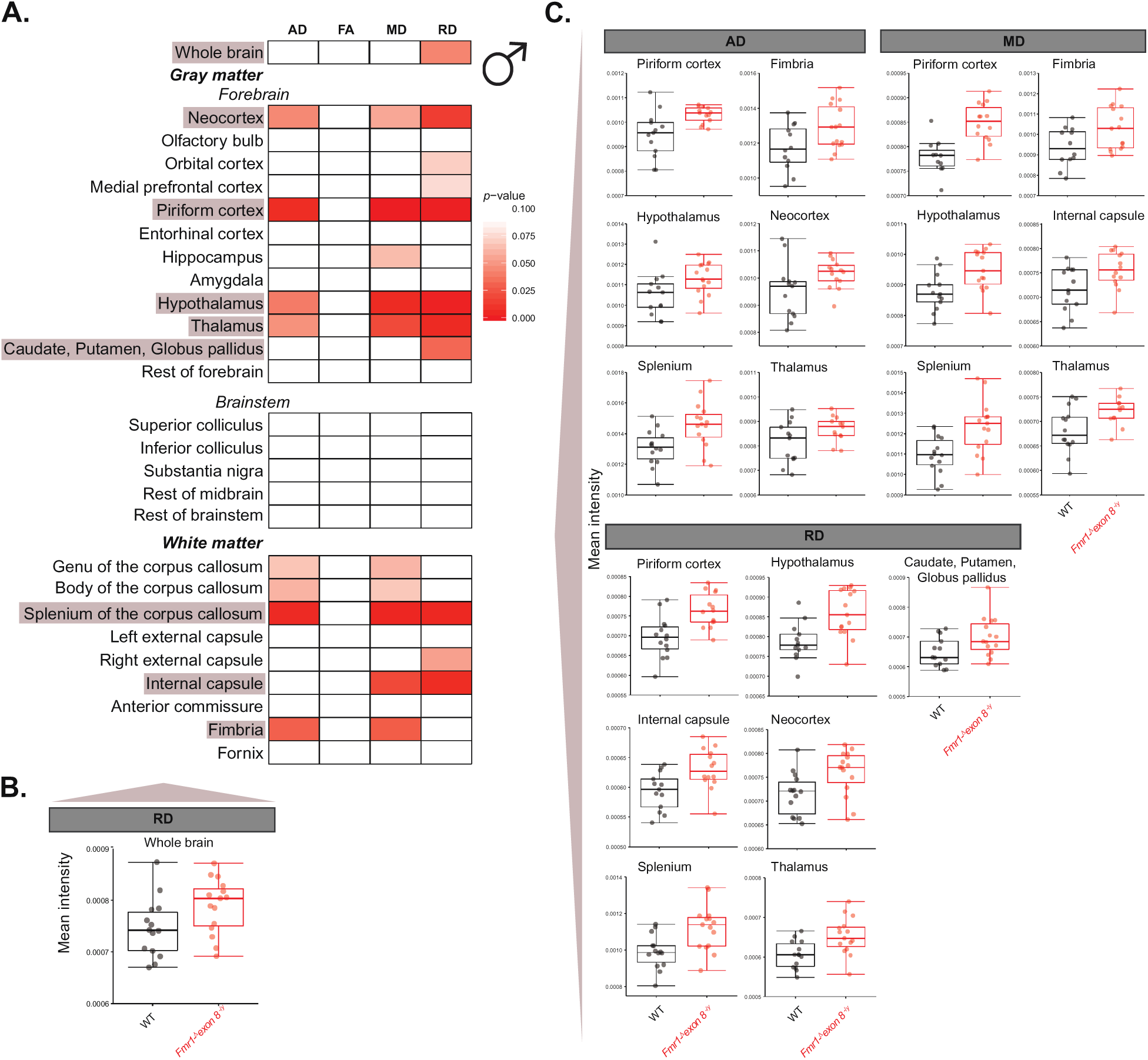
Changes in diffusion indices in *Fmr1-^Δ^exon 8^-/y^* rats and male WT littermates. **(A)** Heatmap of *p*-values from ANOVA and Mann-Whitney U tests for the effect of genotype on AD, FA, MD, and RD using DTI where an increase in red denotes increased significance. **(B)** Boxplot of mean intensity of RD being significantly increased across the whole brain in *Fmr1-^Δ^exon 8^-/y^* rats. **(C)** Boxplots of the brain regions with significantly increased mean intensity of AD, MD, and RD across the multiple brain regions shown in *Fmr1-^Δ^exon 8^-/y^* rats, (WT: N = 14, *Fmr1-^Δ^exon 8^-/y^*: N = 15).

Contrary to the males, when we assessed these parameters in female WT, *Fmr1-^Δ^exon 8^-/-^*, and *Fmr1-^Δ^exon 8^-/+^* littermates, we found that the brainstem was the most affected region. Genotype had a significant effect on AD, MD, and RD in the inferior colliculus (**Figure 4A and B**; ANOVA, *p* = 0.0022, *p* = 0.0061, and *p* = 0.021), substantia nigra (**Figure 4A and B**; ANOVA, *p* = 0.0048, *p* = 0.0057, and *p* = 0.0095), and the rest of the midbrain (**Figure 4A and B**; ANOVA, *p* = 0.29, *p* = 0.0059, and *p* = 0.01). Interestingly, there was a dose-dependent effect of Fmrp-^Δ^KH1 expression where AD, MD, and RD were highest in WT rats, lower in *Fmr1-^Δ^exon 8^-/+^* rats, and lowest in *Fmr1-^Δ^exon 8^-/-^* rats. AD, MD, and RD were all decreased in *Fmr1-^Δ^exon 8^-/-^* compared to WT rats in the inferior colliculus (**Figure 4B**; Tukey HSD, *p adj.* = 0.0017, *p adj.* = 0.0043, and *p adj.* = 0.015); however, only the decrease in AD survived a Bonferroni correction. It was the only finding in the analysis of the females to survive this correction. The substantia nigra was similarly decreased in all three indices in *Fmr1-^Δ^exon 8^-/-^* compared to WT rats (**Figure 4B**; Tukey HSD, *p adj.* = 0.0051, *p adj.* = 0.005, and *p adj.* = 0.0082) and decreased in *Fmr1-^Δ^exon 8^-/-^* compared to *Fmr1-^Δ^exon 8^-/+^* rats in AD and MD (**Figure 4B**; Tukey HSD, *p adj.* = 0.03 and *p adj.* = 0.049). In the rest of the midbrain, which constituted everything in the midbrain that was not otherwise covered by the other specific regions in our analysis, all three indices were also lower in *Fmr1-^Δ^exon 8^-/-^* compared to WT rats (**Figure 4B**; Tukey HSD, *p adj.* = 0.025, *p adj.* = 0.006, and *p adj.* = 0.012). MD and RD were decreased in *Fmr1-^Δ^exon 8^-/-^* compared to *Fmr1-^Δ^exon 8^-/+^* rats (**Figure 4B**; Tukey HSD, *p adj.* = 0.033 and *p adj.* = 0.039). Finally, a significant effect was observed in the amygdala, with increased FA in *Fmr1-^Δ^exon 8^-/-^* compared to *Fmr1-^Δ^exon 8^-/+^* rats (**Figure 4B**; Tukey HSD, *p adj.* = 0.01). This change in FA was not dose-dependent and the difference was largely due to the high variability in the *Fmr1-^Δ^exon 8^-/-^* rats. This difference in FA could be driven by impairments to the myelin within the amygdala or neighboring voxels of white matter. Moreover, all of these effects in these gray matter regions could be driven by deficits in the minimal amount of axonal projections that exists within them or near them, especially in the afferent and efferent projections to these regions. Or, they could be due to gray matter-specific deficits, such as increased spine head density. Notably, while AD, MD, and RD decreased in female *Fmr1-^Δ^exon 8^-/-^* rats compared to WT controls, they increased in the male *Fmr1-^Δ^exon 8^-/y^* rats. This raises the question of whether there would be an interaction between sex and genotype, which can only be addressed by comparing both in one analysis.

**Figure 4.**
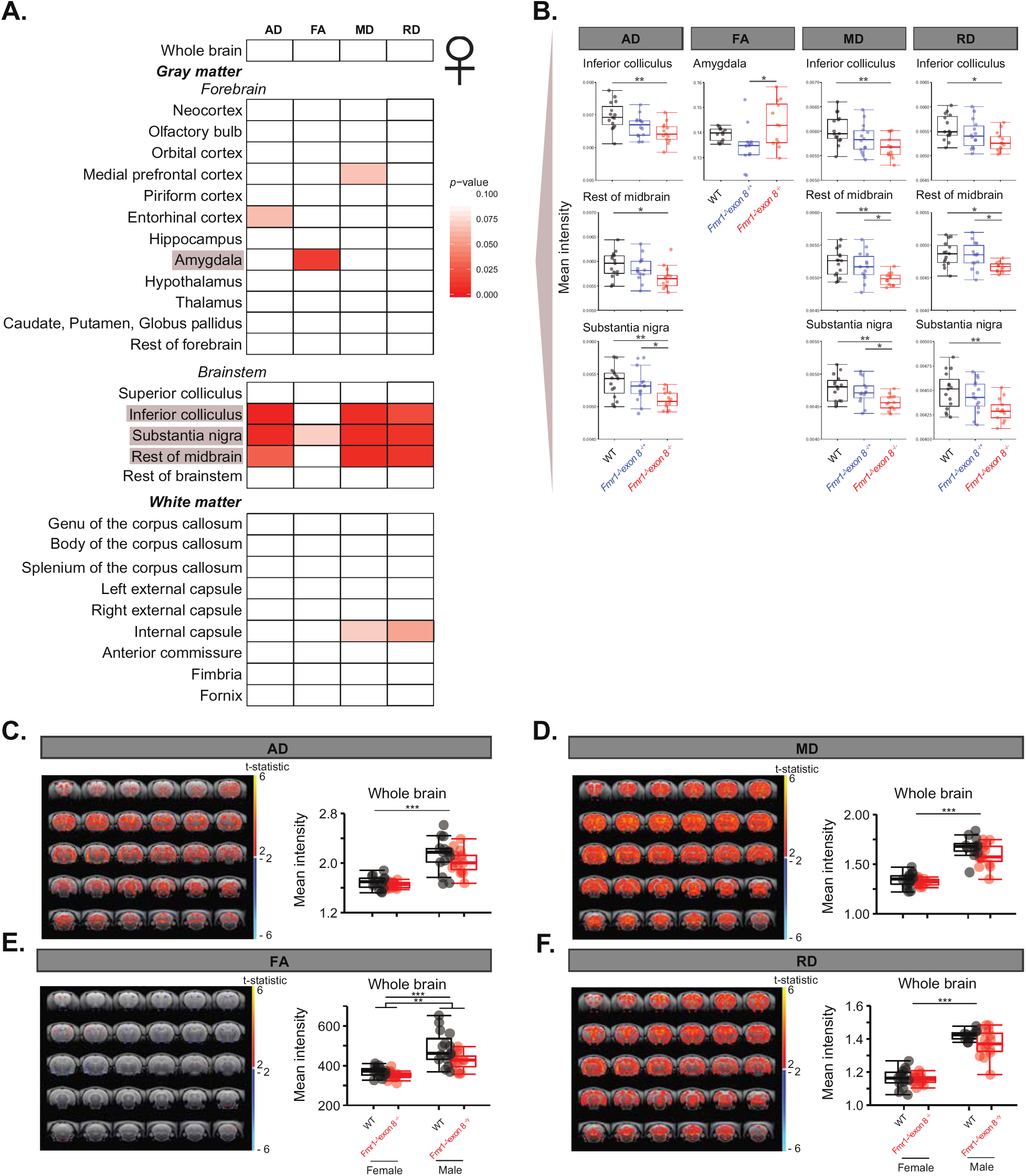
Changes in diffusion indices in DTI images from *Fmr1-^Δ^exon 8^-/-^*, *Fmr1-^Δ^exon 8^-/+^*, and female WT littermates and in whole brain of male and female *Fmr1-^Δ^exon 8* rats and WT littermates. **(A)** Heatmap of *p*-values from ANOVA and Kruskal-Wallis tests for the effect of genotype on AD, FA, MD, and RD where an increase in red denotes increased significance. **(B)** Boxplots of the mean intensity of AD, MD, and RD being significantly decreased in the inferior colliculus, rest of midbrain, and substantia nigra in *Fmr1-^Δ^exon 8^-/-^* rats. Significance bars represent pair-wise comparisons from either a Tukey HSD or Dunn’s test. Maps of the t-statistic across the brain, showing the voxels that are increased or decreased in mean intensity in the male rats compared to the female rats and boxplots of group means across the whole brain for **(C)** AD, **(D)** MD, **(E)** FA, and **(F)** RD where the significance of the pair-wise comparisons from the Tukey HSD or LM is reported, (N = 15/genotype), \*\*\**p* < 0.001, \*\**p* < 0.01, \**p* < 0.05.

### Combining sexes shows effect of genotype and uncovers major sex differences

We validated our findings with the *Fmr1-^Δ^exon 8^-/+^* rats excluded from the analysis using the segmented average from the semi-automated pipeline as a template (see **Supplementary Table 1** for ROIs) and found multiple regions to be altered. FA was reduced in the anterior commissure (**Supplementary Figure 3A**; LM, *p* = 0.00039), neocortex, and rest of forebrain in the *Fmr1-^Δ^exon 8* rats (**Supplementary Figure 3A**; LM, *p* = 0.00086 and *p* = 0.00087, respectively). Additionally, we observed a substantial decrease in MD in the inferior colliculus in both sexes, similar to the decrease we observed specifically in the *Fmr1-^Δ^exon 8^-/-^* rats compared to their WT and *Fmr1-^Δ^exon 8^-/+^* littermates (**Supplementary Figure 3B**; Tukey HSD, *p adj.* = 0.0013). Furthermore, RD was reduced in the amygdala (**Supplementary Figure 3C**; LM, *p* = 0.00095) and hippocampus (**Supplementary Figure 3C**; Tukey HSD, *p adj.* = 0.0015). Unexpectedly, while there were no significant interactions between genotype and sex after correcting for multiple comparisons, there were highly significant effects of sex on these DTI metrics in each region where FA (**Supplementary Figure 3A**; LM, *p* = 0.0029, *p* = 5.25 x 10^-5^, *p* = 7.71 x 10^-4^), MD (**Supplementary Figure 3B**; Tukey HSD, *p adj.* 2.83 x 10^-10^), and RD (**Supplementary Figure 3C**; LM, *p* = 4.04 x 10^-9^ and Tukey HSD, *p adj.* < 1.0 x 10^-20^) were all higher in the males. All nominal *p*-values survived a correction for multiple comparisons, except the *p*-value for the effect of sex on FA in the anterior commissure and genotype on RD in the hippocampus. In fact, across the whole brain, the males had much higher AD (**Figure 4C**; LM, *p* = 7.45 x 10^-12^), MD (**Figure 4D**; Tukey HSD, *p adj.* < 1.0 x 10^-20^), FA (**Figure 4E**; LM, *p* = 5.07 x 10^-9^), and RD (**Figure 4F**; Tukey HSD, *p adj.* < 1.0 x 10^-20^) than females, in which all nominal *p*-values survived a correction for multiple comparisons.

### Axon integrity is disrupted in the splenium of *Fmr1-^Δ^exon 8^-/y^* rats

The splenium stood out in the DTI analysis as a potential locus for impairments in attention because it was increased in AD, MD, and RD in *Fmr1-^Δ^exon 8^-/y^* rats. We examined it further at the level of its ultrastructure using electron microscopy, as differences in diffusion metrics could be reflected by changes at the ultrastructural level. We measured the caliber of the myelinated and unmyelinated axons, estimated the number of myelinated axons in our sample, and determined the thickness of the myelin in *Fmr1-^Δ^exon 8^-/y^* rats and their sex-matched WT littermates. We found that the caliber of myelinated axons was trending to be significantly reduced in *Fmr1-^Δ^exon 8^-/y^* rats (**Figure 5A**; LM, *p* = 0.056). The reduced caliber of these myelinated axons could restrict diffusion along the axis of the axon and underlie the increase in AD seen with DTI in the *Fmr1-^Δ^exon 8^-/y^* rats. Furthermore, while there was no difference in the number of myelinated axons per field (**Figure 5B**; LM, *p* = 0.14), when we multiplied the mean caliber of the myelinated axons by the number represented in a given field, we found that the myelinated axons occupied significantly less of the total field area in *Fmr1-^Δ^exon 8^-/y^* rats (**Figure 5C**; LM, *p* = 0.0012). This suggests that there is more room for diffusion outside of the myelinated axons, which could lead to an increase in MD and RD. However, it is unclear whether MRI is sensitive enough to capture these types of changes. Notably, we did not see deficits in myelin integrity. The thickness of the myelin was roughly equal in male WT rats and *Fmr1-^Δ^exon 8^-/y^* rats (**Figure 5D**; LM, *p* = 0.28). Interestingly, however, the mean g-ratio was reduced in the *Fmr1-^Δ^exon 8^-/y^* rats (**Figure 5E**; LM, *p* = 0.02). To assess whether reduced axonal caliber could be driving this decrease, we examined the relationship between g-ratio and axon caliber. We found a significant interaction between genotype and axon caliber where smaller axons had decreased g-ratios in *Fmr1-^Δ^exon 8^-/y^* rats (**Figure 5F**; LM, *p* = 0.0012). These lower g-ratios in the smaller axons could be due to thicker myelin or reduced internode length. Additionally, thicker myelin surrounding the smaller axons could further restrict diffusion along these axons.

**Figure 5.**
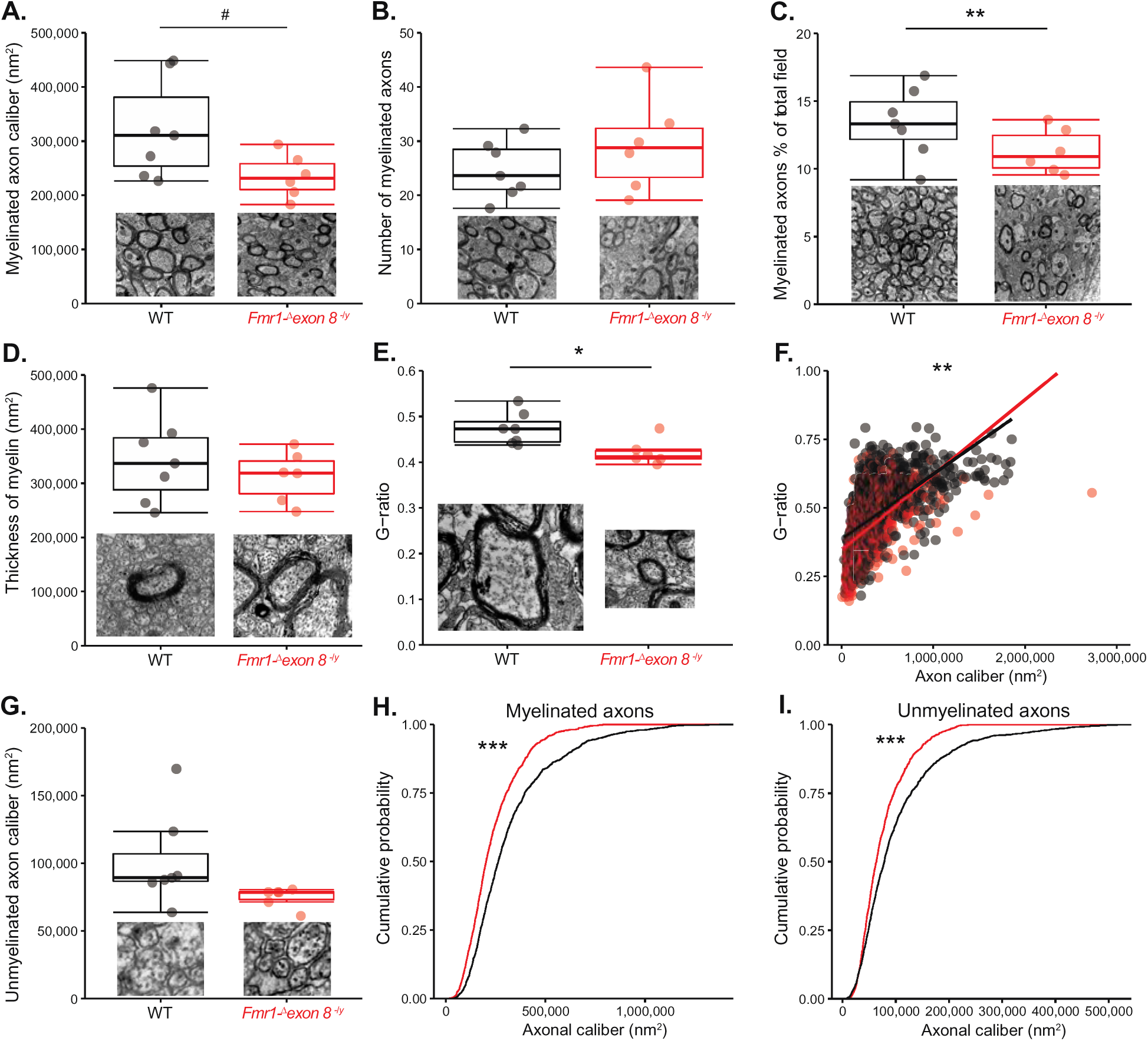
Ultrastructure of myelinated and unmyelinated axons in the splenium of the corpus callosum of *Fmr1-^Δ^exon 8^-/y^* rats and male WT littermates. Boxplots of **(A)** the caliber and **(B)** number of myelinated axons, **(C)** the percentage of the total image that contains myelinated axons, **(D)** the thickness of the myelin, and **(E)** the g-ratio of myelinated axons. **(F)** The g-ratios compared to the caliber of the axons with their linear models. **(G)** Boxplot of axon caliber of unmyelinated axons. Empirical distribution functions of **(H)** myelinated and **(I)** unmyelinated axons (WT: N = 7; *Fmr1-^Δ^exon 8^-/y^*: N = 6), \*\*\**p* < 0.001, \*\**p* < 0.01, \**p* < 0.05, ^#^*p* < 0.1. The insets are examples of the image fields or axons.

Unlike myelinated axons, the caliber of unmyelinated axons was unaffected in the *Fmr1-^Δ^exon 8^-/y^* rats (**Figure 5G**; LM, *p* = 0.1). However, interestingly, when we evaluated the proportion of axons at each caliber, we found a significant difference comparing the cumulative frequencies of axonal calibers in the *Fmr1-^Δ^exon 8^-/y^* rats compared to their WT littermates of both myelinated and unmyelinated axons (**Figure 5H and I**; Kolmogorov-Smirnov test, *p* < 2.2 x 10^-16^ and *p* < 9.78 x 10^-11^, D = 0.19 and D = 0.14). These frequency plots show that the fields from the *Fmr1-^Δ^exon 8^-/y^* rats contained smaller myelinated (< 750,000 nm^2^) and unmyelinated (∼50,000 to ∼250,000 nm^2^) axons and did not contain larger (> 750,000 nm^2^) myelinated and unmyelinated (> 250,000 nm^2^) axons that were present in the WT images. Overall, the impairments in the splenium in *Fmr1* KO mice that were discovered using DTI appear to be at least partially explained by a general reduction in axonal caliber.

## Discussion

In this study, we discovered structural perturbations in several brain regions of *Fmr1-^Δ^exon 8* rats, including in specific domains of white matter, and decreased caliber of axons in the splenium of the corpus callosum. Therefore, a loss of function of the KH1 domain of Fmrp and/or low expression of Fmrp is able to cause large-scale anatomical dysmorphology that may result from reductions in axonal integrity. Importantly, we discovered volumetric changes and alterations in diffusion in both gray and white matter regions that are also impaired in *Fmr1* KO mice and individuals with FXS.

Using a semi-automated region-based analysis of T2 images, we determined that the superior colliculus increased in absolute volume in *Fmr1-^Δ^exon 8^-/y^* rats and validated this in a fully automated analysis. The superior colliculus is also impaired in *Fmr1* KO mice [6] and implicated in the orienting component of attention. Second, the genu of the corpus callosum increased in both absolute and relative volume in *Fmr1-^Δ^exon 8^-/+^* rats compared to WT littermates. The genu of the corpus callosum is increased in overall diffusion in girls with FXS [4]. Third, the olfactory bulb increased in absolute volume in *Fmr1-^Δ^exon 8^-/+^* rats. Fmrp expression is relatively high in the olfactory bulb compared to the rest of the brain in *Fmr1* KO mice, making it a vulnerable region to changes in Fmrp expression [22]. Lastly, the hypothalamus decreased in relative volume in *Fmr1-^Δ^exon 8^-/-^* rats. The hypothalamus is also decreased in volume in the most comprehensive MRI study thus far that used a mouse model of FXS, the *Fmr1* KO on the FVB background strain [5]. Unlike these rodent models of FXS, it is enlarged in boys with FXS [23]. It is thought that the abnormal volume of the hypothalamus in FXS could be related to their dysfunctional endocrine system that causes macroorchidism, diminished pubertal growth, and elevated levels of cortisol in basal conditions and in response to stress [24]. Disturbances in the neuroendocrine system have also been identified in *Fmr1* KO mice, which have similarly elevated basal glucocorticoid hormones and a heightened response to stress [25]. The fully automated analysis further identified changes in the volumes of the fourth and lateral ventricles. The decrease in volume of the fourth ventricle could be due to the increase in size of the PAG and the increase in volume of the lateral ventricle could be due to subtle changes in volume of the brain regions surrounding it.

While a region-based approach is sensitive to volumetric changes at the anatomical level, we also applied a voxel-based approach further to probe changes at a smaller scale. Using this voxel-based approach, we found the PAG to be increased in both *Fmr1-^Δ^exon 8^-/y^* and *Fmr1-^Δ^exon 8^-/-^* rats. The PAG is also increased in volume in *Fmr1* KO mice on the FVB background strain [5]. Notably, the PAG has not garnered much attention in studies of individuals with FXS and there are no reports of its volume in this population. Because it is enlarged in both a mouse and rat model of FXS, crucially in both sexes of a rat model, its volume deserves consideration in individuals with FXS.

In our primary semi-automated region-based analysis of DTI images, we discovered alterations in diffusion in both gray and white matter regions that are also impaired in *Fmr1* KO mice and individuals with FXS. In the forebrain, the caudate/putamen and internal capsule are similarly smaller and the fimbria and inferior colliculus are similarly enlarged in the *Fmr1* KO mouse on the FVB background strain compared to WT controls [5]. This reduction in size of the caudate nucleus is opposite of what is often documented in individuals with FXS, but the caudate and putamen are separate nuclei in humans, making them difficult to compare to their rodent homolog. Unlike the caudate nucleus, the fimbria and inferior colliculus have been unexplored as regions that could underlie FXS pathology. While the internal capsule also changed in the opposite direction to what is documented in humans with FXS and in *Fmr1* KO mice on the FVB background strain, it has increased MD and RD in its anterior limbs [4], which complements what we found in *Fmr1-^Δ^exon 8^-/y^* rats. Further complementing our results, the splenium and thalamus show increased AD, MD, and RD in girls with FXS [4], the thalamus has increased gray matter volume [23], and FA is decreased in the splenium of *Fmr1* KO mice [7]. Interestingly, however, no prior studies have examined the anatomy of the substantia nigra in FXS even though dopaminergic tone is dampened [26], likely because this dysfunction is most often attributed to deficits in the frontostriatal pathway. An increase in MD in the substantia nigra typically indicates a loss of dopaminergic neurons in Parkinson’s disease [27]. Interestingly, *Fmr1* KO mice have fewer tyrosine hydroxylase-expressing neurons in the substantia nigra [28]. The decrease in diffusion that we observed could be related to the integrity of the dopaminergic neurons in *Fmr1-^Δ^exon 8^-/-^* rats. All of these anatomical perturbations could be related to the deficits we see in behavior and transcription of *Fmr1-^Δ^exon 8* rats [11]. This is worth further exploration.

When we combined the males and females in one analysis, we discovered altered diffusion in the neocortex and inferior colliculus of *Fmr1-^Δ^exon 8* rats, which were both impaired in male and female rats when they were individually compared to their sex-matched controls. Additionally, we found that FA in the anterior commissure and forebrain and RD in the amygdala and hippocampus were decreased in *Fmr1-^Δ^exon 8* rats. The finding that diffusivity in the anterior commissure is impaired further suggests that this *Fmr1-exon 8* deletion can lead to white matter impairments. Interestingly, we discovered a general increase in diffusion across the board in males compared to females, a potential explanation for the observed increase in diffusion in the male *Fmr1-^Δ^exon 8* rats and decrease in the female *Fmr1-^Δ^exon 8* rats compared to WT controls. A similar trend was also identified for FA and AD in humans [29]. These authors attributed this sex difference to hormonal exposure. Furthermore, it has been shown that males have greater within-hemisphere connectivity and females have greater between-hemisphere connectivity, which could potentially affect these measures [30]. This sex difference in rats should be replicated in another strain.

The splenium showed increases in all three measures of diffusion. Because the splenium is related to attention and this finding recapitulates what is seen in girls with FXS, we decided to explore its integrity further at the level of histology. Using electron microscopy, we found that axons were decreased in caliber, myelinated axons occupied less of the image, and smaller axons had thicker myelin in *Fmr1-^Δ^exon 8^-/y^* rats. It is possible that the reduced caliber of these axons and the increased thickness of myelin surrounding smaller axons could restrict diffusion along the axis of the axon, contributing to the increase in AD we identified with DTI. It is further possible that because the myelinated axons inhabited less of the image, there could be more room for MD and RD. This association warrants further study. Regardless of the relationship, this is the first time that axon caliber was assessed in the adult and in the corpus callosum of a model of FXS. Axons in the cerebellum of *Fmr1* KO mice at postnatal days 7 and 15 have been assessed, but no difference in axon diameter was observed [31]. This raises the question whether reduction in axonal caliber is specific to the splenium. Axonal caliber can depend upon the amount of energy available, which is mostly regulated by mitochondrial function [32]. In the immature neurons of the dentate gyrus of adult *Fmr1* KO mice, mitochondria are shorter and mitochondria-related genes show decreased expression [33]. Furthermore, mitochondrial membrane potential in cultured neurons is reduced and oxidative stress is increased, indicating mitochondrial dysfunction. It should be assessed whether the same phenomena could be occurring in the axons of the splenium of *Fmr1-^Δ^exon 8^-/y^* rats, which could offer a potential mechanism for the reduction in axonal caliber we identified. Interestingly, we previously found that PGC-1α (also known as PPARGC1A), a master regulator of mitochondria biogenesis that is also an Fmrp target, is decreased in expression in *Fmr1-^Δ^exon 8^-/y^* rats [11]. It would be worthwhile to determine whether PGC-1α plays a role in this deficit in axonal caliber. Weakening of the cytoskeleton could also cause a reduction in caliber. Fmrp regulates the translation of mRNAs that encode for proteins that regulate cytoskeleton remodeling. Because Fmrp localizes to axons and associates with translational machinery in rats [34], it is possible that the expression of proteins that regulate the integrity of the cytoskeleton is dysregulated in *Fmr1-^Δ^exon 8^-/y^* rats, leading to malformed axonal cytoskeleton.

The brain regions we found to be altered that are also impaired in FXS and involved in attention networks are worth further exploration as targets for treatment of attention deficits in FXS because they are also implicated in ADHD. For example, in ADHD, MD in the caudate, putamen, and thalamus is correlated with increased reaction time on a flanker test [35]. Furthermore, the internal capsule has impaired white matter integrity [36]. Lastly, there is higher MD and RD, and lower FA [37] in the splenium. The lower FA phenotype is especially prominent in patients with the inattention phenotype [36], the specific deficit we discovered in the *Fmr1-^Δ^exon 8* rats [14]. Therefore, the relationship between the altered structure of these regions and attentional functioning in FXS models should be studied more directly in the same animals.

Sensory deficits have also been replicated in FXS models [38]. Many of the brain regions that were perturbed in our study are implicated in sensory perception. To begin with, three regions that are involved in olfaction, the piriform cortex, olfactory bulb, and anterior commissure were altered in *Fmr1-^Δ^exon 8* rats. Olfaction has not been well characterized in FXS, but it is impaired in rodent models of FXS [39]. Individuals with FXS can also be hypersensitive to auditory and visual stimuli. We found that the inferior and superior colliculus, which are in the auditory and visual pathways, respectively, are altered in *Fmr1-^Δ^exon 8* rats. Additionally, we identified alterations in the PAG, which, among other functions, is involved in somatosensory perception and the thalamus, which relays sensory information to the cortex, to be increased in diffusion. C-fos expression in the PAG and thalamus is excessive after an auditory stimulus that causes audiogenic-induced seizures in *Fmr1* KO mice [40, 41]. The role that these regions play in hypersensitivity in FXS has not garnered much attention. The connection between the structural alterations we identified and the sensory deficits that are often observed in FXS models should be further explored.

In summary, we have shown here that a specific deletion of exon 8 of the *Fmr1* gene is sufficient to cause FXS-like changes in neuroanatomy in rat, specifically in the axons of a major pathway. Therefore, white matter in FXS deserves further examination. The regions we found to be disrupted could serve as potential cross-species non-invasive biomarkers for FXS.

## Acknowledgments

We thank Emma Huang, Jenni Li, Olamide Olawuni, Aziza Ndaw, Sebastian Sukdeo, Claudia Yang, Alice Cheng, and Eilam Doron, who contributed to this work, and The Mount Sinai Microscopy Core and BioMedical Molecular Imaging Institute for their services in carrying out the electron microscopy and magnetic resonance imaging. This work was supported by the Seaver Foundation (to JDB, HHN, and CEMG); Autism Speaks (to JDB); and the National Institute of Mental Health (grant number F31 MH115656 to CEMG). Carla E. M. Golden was a Seaver Graduate Fellow at the time of this study.

## Conflict of Interest

There are no conflicts of interest.

Supplementary information is available at TP’s website.

---

**Supplementary Figure 1.**
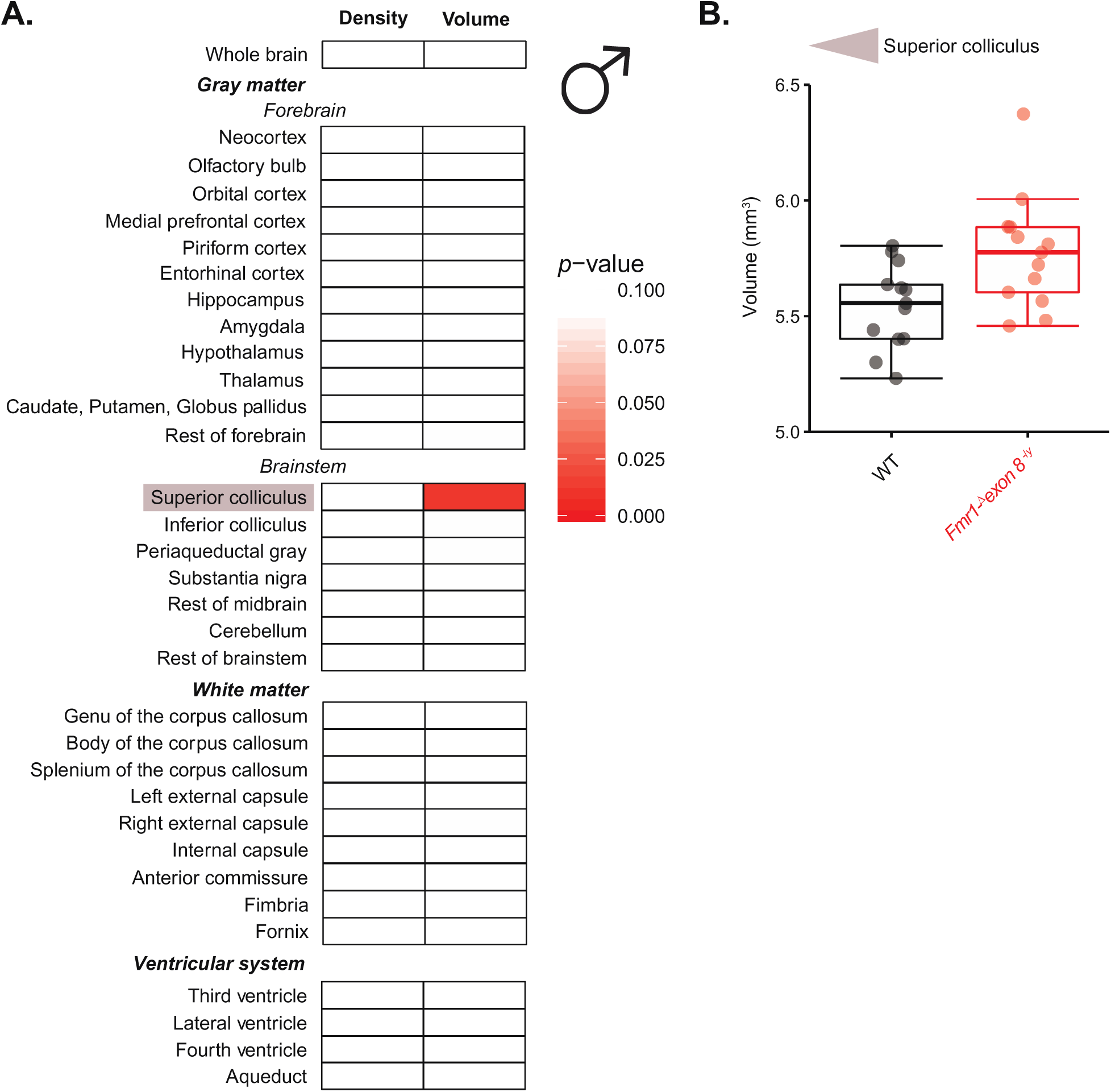
Volumes and tissue density of brain regions of *Fmr1-^Δ^exon 8^-/y^* rats and male WT littermates. **(A)** Heatmap of *p*-values from ANOVA and Mann-Whitney U tests for the effect of genotype on volume and mean intensity in T2 images where an increase in red denotes increased significance. **(B)** Boxplot of the significantly increased volume of the superior colliculus, (WT: N = 15; *Fmr1-^Δ^exon 8^-/y^*: N = 14).

**Supplementary Figure 2.**
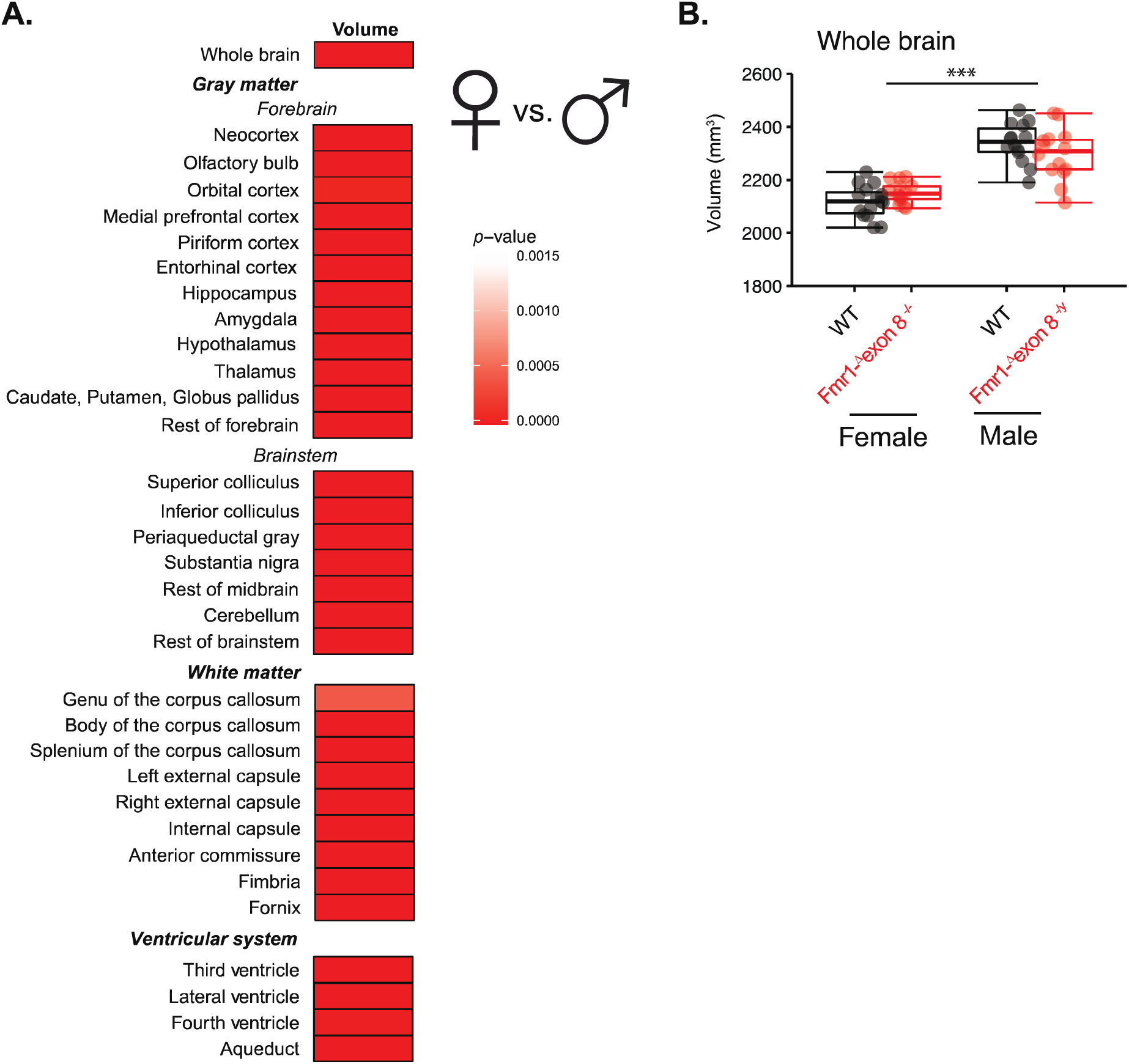
Volumes of brain regions of male and female *Fmr1-^Δ^exon 8* rats and WT littermates. **(A)** Heatmap of *p*-values from ANOVA and LM tests for the effect of sex on volume in T2 images where an increase in red denotes increased Bonferroni-corrected significance. **(B)** Boxplot of the volume of the whole brain, (N = 15/group), \*\*\**p* < 0.001.

**Supplementary Figure 3.**
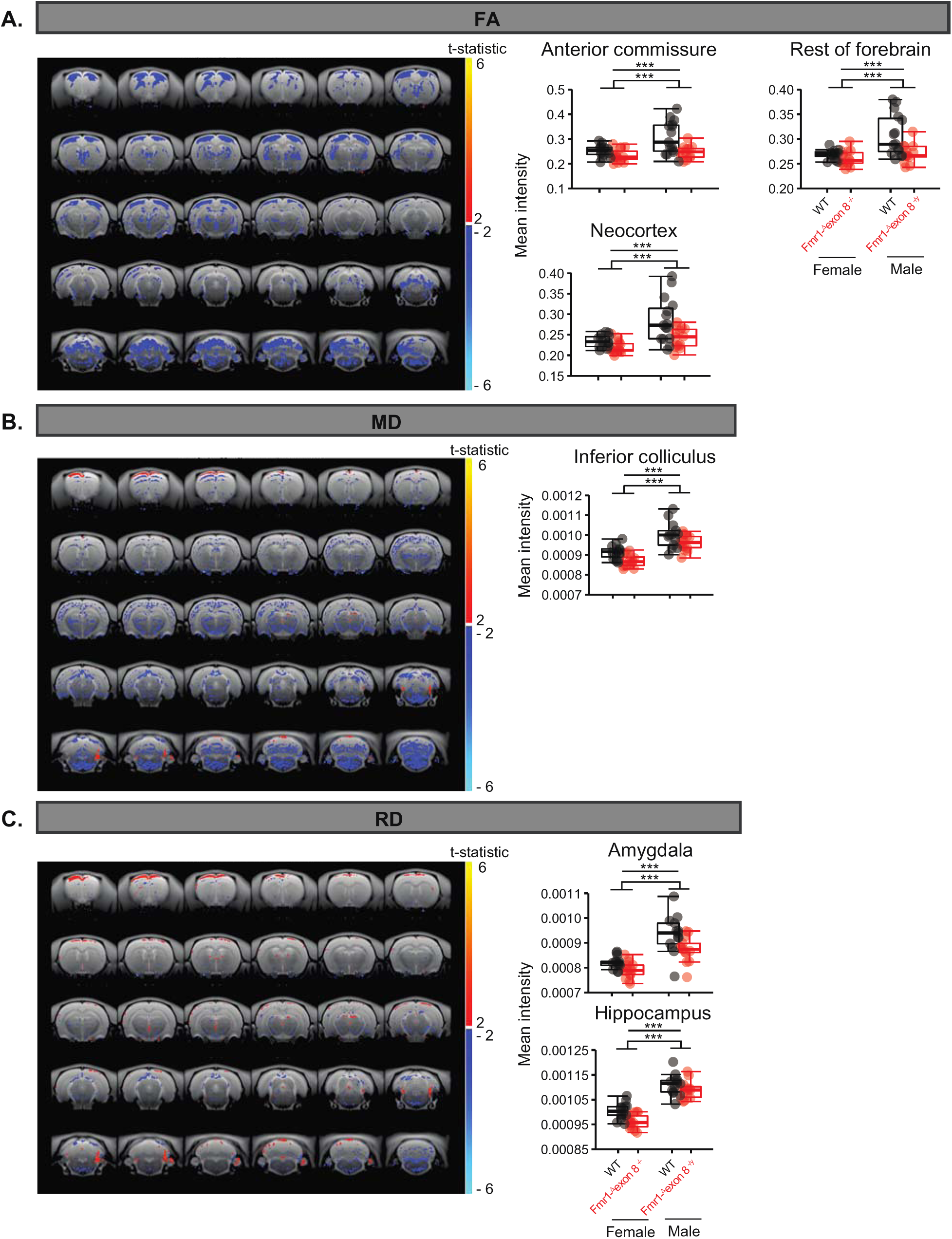
Voxel-wise analysis of DTI images from *Fmr1-^Δ^exon 8^-/-^* and *Fmr1-^Δ^exon 8^-/y^* rats and WT littermates. Maps of the t-statistic across the brain, showing the voxels that are increased or decreased in mean intensity in the *Fmr1-^Δ^exon* rats compared to WT rats and boxplots of group means for **(A)** FA, **(B)** MD, and **(C)** RD where the significance of the pair-wise comparisons from the Tukey HSD or LM is reported, (N = 15/genotype), \*\*\**p* < 0.001.

## References

1. Bagni, C., F. Tassone, G. Neri, and R. Hagerman, Fragile X syndrome: causes, diagnosis, mechanisms, and therapeutics. J Clin Invest, 2012. 122(12): p. 4314–22.

2. Hagerman, R.J., et al., Fragile X syndrome. Nat Rev Dis Primers, 2017. 3: p. 17065.

3. Bray, S., et al., Aberrant frontal lobe maturation in adolescents with fragile X syndrome is related to delayed cognitive maturation. Biol Psychiatry, 2011. 70(9): p. 852–8.

4. Villalon-Reina, J., et al., White matter microstructural abnormalities in girls with chromosome 22q11.2 deletion syndrome, Fragile X or Turner syndrome as evidenced by diffusion tensor imaging. Neuroimage, 2013. 81: p. 441–454.

5. Lai, J.K., J.P. Lerch, L.C. Doering, J.A. Foster, and J. Ellegood, Regional brain volumes changes in adult male FMR1-KO mouse on the FVB strain. Neuroscience, 2016. 318: p. 12–21.

6. Kay, R.B., N.A. Gabreski, and J.W. Triplett, Visual subcircuit-specific dysfunction and input-specific mispatterning in the superior colliculus of fragile X mice. J Neurodev Disord, 2018. 10(1): p. 23.

7. Haberl, M.G., et al., Structural-functional connectivity deficits of neocortical circuits in the Fmr1 (-/y) mouse model of autism. Sci Adv, 2015. 1(10): p. e1500775.

8. Hall, S.S., H. Jiang, A.L. Reiss, and M.D. Greicius, Identifying large-scale brain networks in fragile X syndrome. JAMA Psychiatry, 2013. 70(11): p. 1215–23.

9. Zerbi, V., et al., Inhibiting mGluR5 activity by AFQ056/Mavoglurant rescues circuit-specific functional connectivity in Fmr1 knockout mice. Neuroimage, 2019. 191: p. 392–402.

10. Richter, J.D., G.J. Bassell, and E. Klann, Dysregulation and restoration of translational homeostasis in fragile X syndrome. Nat Rev Neurosci, 2015. 16(10): p. 595–605.

11. Golden, C.E.M., et al., Deletion of the KH1 Domain of Fmr1 Leads to Transcriptional Alterations and Attentional Deficits in Rats. Cereb Cortex, 2019. 29(5): p. 2228–2244.

12. Engineer, C.T., et al., Degraded speech sound processing in a rat model of fragile X syndrome. Brain Res, 2014. 1564: p. 72–84.

13. Radyushkin, K., et al., Neuroligin-3-deficient mice: model of a monogenic heritable form of autism with an olfactory deficit. Genes Brain Behav, 2009. 8(4): p. 416–25.

14. Golden, C.E.M., et al., Deletion of the KH1 Domain of Fmr1 Leads to Transcriptional Alterations and Attentional Deficits in Rats. Cereb Cortex, 2019.

15. Budin, F., et al., Fully automated rodent brain MR image processing pipeline on a Midas server: from acquired images to region-based statistics. Front Neuroinform, 2013. 7: p. 15.

16. Lerch, J.P., J.G. Sled, and R.M. Henkelman, MRI phenotyping of genetically altered mice. Methods Mol Biol, 2011. 711: p. 349–61.

17. Avants, B.B., et al., The Insight ToolKit image registration framework. Front Neuroinform, 2014. 8: p. 44.

18. Winklewski, P.J., et al., Understanding the Physiopathology Behind Axial and Radial Diffusivity Changes-What Do We Know? Front Neurol, 2018. 9: p. 92.

19. Spring, S., J.P. Lerch, and R.M. Henkelman, Sexual dimorphism revealed in the structure of the mouse brain using three-dimensional magnetic resonance imaging. Neuroimage, 2007. 35(4): p. 1424–33.

20. Krauzlis, R.J., L.P. Lovejoy, and A. Zenon, Superior colliculus and visual spatial attention. Annu Rev Neurosci, 2013. 36: p. 165–82.

21. Tolomeo, S., S. Gray, K. Matthews, J.D. Steele, and A. Baldacchino, Multifaceted impairments in impulsivity and brain structural abnormalities in opioid dependence and abstinence. Psychol Med, 2016. 46(13): p. 2841–53.

22. Zorio, D.A., C.M. Jackson, Y. Liu, E.W. Rubel, and Y. Wang, Cellular distribution of the fragile X mental retardation protein in the mouse brain. J Comp Neurol, 2017. 525(4): p. 818–849.

23. Hoeft, F., et al., Morphometric spatial patterns differentiating boys with fragile X syndrome, typically developing boys, and developmentally delayed boys aged 1 to 3 years. Arch Gen Psychiatry, 2008. 65(9): p. 1087–97.

24. Hessl, D., S.M. Rivera, and A.L. Reiss, The neuroanatomy and neuroendocrinology of fragile X syndrome. Ment Retard Dev Disabil Res Rev, 2004. 10(1): p. 17–24.

25. Lauterborn, J.C., Stress induced changes in cortical and hypothalamic c-fos expression are altered in fragile X mutant mice. Brain Res Mol Brain Res, 2004. 131(1-2): p. 101–9.

26. Hagerman, R., J. Lauterborn, J. Au, and E. Berry-Kravis, Fragile X syndrome and targeted treatment trials. Results Probl Cell Differ, 2012. 54: p. 297–335.

27. Ofori, E., et al., Free water improves detection of changes in the substantia nigra in parkinsonism: A multisite study. Mov Disord, 2017. 32(10): p. 1457–1464.

28. Paul, K., D.V. Venkitaramani, and C.L. Cox, Dampened dopamine-mediated neuromodulation in prefrontal cortex of fragile X mice. J Physiol, 2013. 591(4): p. 1133–43.

29. van Hemmen, J., et al., Sex Differences in White Matter Microstructure in the Human Brain Predominantly Reflect Differences in Sex Hormone Exposure. Cereb Cortex, 2017. 27(5): p. 2994–3001.

30. Ingalhalikar, M., et al., Sex differences in the structural connectome of the human brain. Proc Natl Acad Sci U S A, 2014. 111(2): p. 823–8.

31. Pacey, L.K., et al., Delayed myelination in a mouse model of fragile X syndrome. Hum Mol Genet, 2013. 22(19): p. 3920–30.

32. Perge, J.A., J.E. Niven, E. Mugnaini, V. Balasubramanian, and P. Sterling, Why do axons differ in caliber? J Neurosci, 2012. 32(2): p. 626–38.

33. Shen, M., et al., Reduced mitochondrial fusion and Huntingtin levels contribute to impaired dendritic maturation and behavioral deficits in Fmr1-mutant mice. Nat Neurosci, 2019. 22(3): p. 386–400.

34. Akins, M.R., et al., Axonal ribosomes and mRNAs associate with fragile X granules in adult rodent and human brains. Hum Mol Genet, 2017. 26(1): p. 192–209.

35. Fall, S., L. Querne, A.G. Le Moing, and P. Berquin, Individual differences in subcortical microstructure organization reflect reaction time performances during a flanker task: a diffusion tensor imaging study in children with and without ADHD. Psychiatry Res, 2015. 233(1): p. 50–6.

36. Qiu, M.G., et al., Changes of brain structure and function in ADHD children. Brain Topogr, 2011. 24(3-4): p. 243–52.

37. Chen, L., et al., A systematic review and meta-analysis of tract-based spatial statistics studies regarding attention-deficit/hyperactivity disorder. Neurosci Biobehav Rev, 2016. 68: p. 838–847.

38. Golden, C.E., J.D. Buxbaum, and S. De Rubeis, Disrupted circuits in mouse models of autism spectrum disorder and intellectual disability. Curr Opin Neurobiol, 2018. 48: p. 106–112.

39. Bodaleo, F., C. Tapia-Monsalves, C. Cea-Del Rio, C. Gonzalez-Billault, and A. Nunez-Parra, Structural and Functional Abnormalities in the Olfactory System of Fragile X Syndrome Models. Front Mol Neurosci, 2019. 12: p. 135.

40. Goebel-Goody, S.M., et al., Genetic manipulation of STEP reverses behavioral abnormalities in a fragile X syndrome mouse model. Genes Brain Behav, 2012. 11(5): p. 586–600.

41. Chen, L. and M. Toth, Fragile X mice develop sensory hyperreactivity to auditory stimuli. Neuroscience, 2001. 103(4): p. 1043–50.

